# Making Good Choices: Social Interaction in Mice Mitigates Chronic Stress-Induced Adaptive Changes in Decision Making

**DOI:** 10.1101/2020.03.02.973156

**Authors:** Arish Mudra Rakshasa, Michelle T. Tong

## Abstract

Chronic stress can impact decision-making and lead to a preference for immediate rewards rather than long-term payoffs. Factors that may mitigate these effects of chronic stress on decision-making are under-explored. Here we used a mouse model to investigate the changes in decision-making caused by the experience of chronic stress and the role of social interaction in attenuating these changes. To test decision-making, mice were trained to perform a Cost-Benefit Conflict (CBC) task on a T-maze, in which they could choose between a high-reward, high-risk alternative and a low-reward, low-risk alternative. Mice were either housed in groups or alone throughout the experiment. Both groups of mice underwent a seven-day period of repeated immobilisation to induce chronic stress. Stress levels were confirmed using behavioural (open field test) and physiological (urine corticosterone ELISA) measures. We found a significant increase in frequency of high-risk decisions after exposure to chronic stress among both socially- and individually-housed mice. Crucially, socially-housed mice showed a significantly smaller increase in high-risk decision-making compared to singly-housed mice. These findings suggest that although chronic stress leads to an increase in high-risk decision-making in mice, access to social interaction may mitigate this stress effect.

## Introduction

“Poverty to a large extent is also a state of mind,” said Housing and Urban Development Secretary Ben Carson, a top US government official tasked with increasing access to affordable housing, in a radio interview for National Public Radio [17]. Secretary Carson’s comments reflect a widely held belief that poverty is caused by individual decision-making, where people living in poverty are stereotyped as making bad choices like buying iPhones rather than investing in healthcare [7]. Such comments suggest that people living in poverty make decisions based on inherently flawed values, and would escape the harsh conditions of poverty if only they changed their “state of mind.” Put another way, poor decision-making causes poverty.

This narrative; however, does not account for the effects of poverty-related chronic stress on individuals’ physical and mental health. Defined as prolonged and constant exposure to negative emotional experiences, chronic stress has been linked to negative health outcomes such as increased incidences of coronary heart disease [54], the acceleration of memory impairments [26], and increased tumour growth [51]. In addition, prolonged exposure to stressful environments may be correlated with increased vulnerability to mental health issues such as depression [9, 43] and substance abuse disorders [5, 49]. Other mental health disorders, including post-traumatic stress disorder (PTSD) [2] and schizophrenia [55], may also be triggered or aggravated by chronic stress. Therefore, chronic stress has well-documented negative effects on both physical and mental health of individuals.

Of particular relevance to this study, chronic stress has been shown to affect cognitive processes such as decision-making [8, 25]. More specifically, persistent exposure to stressful situations can lead to increased risk-taking [61]. For instance, military students under severe chronic stress were more likely to fire at targets even when they found that the targets were humans and not humanoid dummies [30]. Chronic stress can also lead to changes in decision-making about personal health, causing individuals to prefer high, immediate rewards rather than delayed payoffs in both food choice [39] and substance use [18]. Mercer and colleagues [33] found that chronically-stressed individuals were more likely to choose “comfort foods” over healthy food. These studies suggest a strong relationship between chronic stress and an increased preference for immediate rewards in decision-making.

A direct causal relationship between chronic stress and decision-making, however, can be challenging to establish for human participants, since human studies necessarily rely on self-reports from participants who are already chronically stressed. In this regard, animal models can provide an effective means of studying the effects of chronic stress on decision-making. Chronic stress may be reliably induced in rodents using several well-established stress protocols [32, 36]. As such, researchers can manipulate physiological and environmental conditions to explore causality of chronic stress effects and assess these effects at different time points. Animal models also allow researchers to better understand biochemical and neural effects of chronic stress. Moreover, rodents that are modified to mimic certain physiological or psychological conditions relevant to humans may provide insight into stress-related responses in humans [50].

Several studies of chronic stress with rodent models [44, 60] have shown that mice exposed to chronic stressors exhibit decreased evaluation of choices and reduced hesitation in making a decision [42]. Rats subjected to such chronic stress make decisions based on habit rather than an evaluation of consequence [14]. Friedman and colleagues [21] have recently demonstrated that mice engage in “aberrant” decision-making behaviour after being subjected to chronic immobilisation stress. When the researchers presented mice with the decision to choose between a high-reward, high-risk alternative and a low-reward, low-risk alternative, they found that chronically-stressed mice “were significantly more likely to choose high-risk/high-reward options than matched controls” [21, p.1192], indicating that mice exposed to chronic stress preferred immediate high payoffs in their decisions, whereas unstressed mice consistently preferred low-risk alternatives. These studies indicate that chronic stress has a significant influence on decision-making, such that it may actually be said to cause individuals to favour high-risk, high-reward decisions. Thus, we recommend a shift from framing high-risk decisions as “aberrant” or “bad,” and suggest, instead, that choosing risky immediate rewards may in fact be “adaptive” in stressful, unpredictable environments.

These results have strong implications for our understanding of human decision-making. They suggest that chronic stress causes a cognitive bias toward high-risk, high-reward choices, which may at times perpetuate conditions of chronic stress. Thus, it is critical to explore factors that may counteract these cyclical effects. There is some evidence that social interaction could attenuate the effects of chronic stress [13]. Regular, healthy social interaction may create a buffer against chronic stress [10], such that the effects of chronic stress may be mediated by the access to social support offered by one’s living conditions [15, 45]. People who do not have access to healthy social interaction, such as older adults living in isolation, seem to face increased stress and risk of cardiovascular disease [58]. Conversely, older adults who receive emotional support from social settings such as churches report lesser effect of financial stressors on self-rated health [28]. Thus, housing conditions and availability of healthy social support may be significant in mitigating the effects of chronic stress.

Given that mice and rats are also social animals, and that responses such as pain sensitivity are modulated by social factors in mice [29], housing conditions among rodent models offer an operationalisation of social support in humans. Social housing was shown to lessen the effects of chronic stress on behaviours such as sexual activity in rats, whereas rats raised in isolation were more vulnerable to chronic stress-induced behavioural changes such as decreased sexual activity [41]. Rats housed socially performed much better on spatial memory tests than those raised in isolation [37], suggesting that social interaction may improve or preserve basic cognitive functions. Additionally, [3] have shown that being subjected to social deprivation represents a stressful condition for mice, and that mice in different housing conditions may use different adaptive responses to these stressors. Notably, social housing in male rats particularly seems to reduce the effects of chronic stress and prevent significant decreases in neurogenesis [56], which may suggest that housing condition may interact significantly with the effects of chronic stress on cognitive functions such as decision-making. Thus, social interaction may be a potent factor in attenuating the effects of chronic stress in both animals and humans.

The current study investigated the possible role of social interaction in attenuating stress-induced changes in decision-making in mice. Although the effects of chronic stress and social isolation on decision-making have separately been studied, to our knowledge, the interaction between these three factors has not yet been explored together. Decision-making and stress were assessed in socially-housed and singly-housed mice using behavioural and physiological measures, with adaptive decision-making defined as a greater tendency to choose a high-reward alternative in conditions of stress. There were two research hypotheses for the current study: (1) Chronic-stress through repeated immobilisation would cause increased high-risk/high-reward decision-making in all mice, and (2) social housing would mitigate the effect of chronic stress on decision-making, such that socially-housed mice would show a smaller increase in high-risk/high-reward decisions than singly-housed mice after chronic stress.

## Methods

### Animals

Thirty male C57BL/6J mice (Charles River Laboratories, 7-8 weeks old at the time of arrival) were used in this experiment. Immediately after arrival at the housing facilities, the mice were randomly assigned to home cages alone (singly-housed, *n* = 15) or in groups of five (socially-housed, *n* = 15). Animals were kept on a 12:12 hour reverse light/dark cycle for the duration of the experiment. All mice were housed at constant temperature (70°F) in standard Plexiglass cages (28 cm x 17 cm x 12 cm) and were given unrestricted access to food and water throughout the study. All procedures performed with rodents were approved by the Institutional Animal Care and Use Committee at Earlham College prior to the start of the study and were conducted in accordance with the U.S. National Research Council Guide for the Care and Use of Laboratory Animals.

### Procedure

This study employed a Pre-Post-Control mixed-measures design to compare effects of chronic stress on decision-making between socially-housed and isolated mice. Briefly, all mice underwent an initial Open Field Test (behavioural measure of stress), urine collection for corticosterone ELISA assays (physiological marker of stress), and the Cost-Benefit Conflict task (measure of decision-making). Following these pre-stress tests, all mice were subjected to seven days of repeated immobilisation to induce chronic stress. Finally, mice were tested again using the open field test, urine corticosterone ELISA assay, and the CBC task to measure post-stress levels of stress and decision-making.

#### Habituation and Pretraining

All mice were first habituated to the experimenter and to the T-maze apparatus (Maze Engineers, Boston, MA; stem 30 cm x 10 cm, arm length 30 cm each) used for the CBC test. Mice were simply allowed to roam on the T-maze for two minutes a day for one week. For pretraining, each maze arm was fitted with a food well (1 cm in diameter) and a clip-on lamp at the far end. Chocolate whole milk purchased from a local grocery store, either pure or diluted with water to an 80% dilution by volume, was used as a reward for mice in the food wells. Bright (1600 lumens) and dim (100 lumens) lights (General Electric LED bulbs) shining on respective arms of the T-maze were used as severe and moderate aversive stimuli, respectively. All mice were trained in receiving the chocolate milk reward from the food wells in the T-maze using 10 repeated forced-choice trials. Habituation and pretraining occurred for five days, during which time all mice were already seperated into singly-housed and socially-housed groups.

#### Open Field Test

All mice were tested in the Open Field Test twice, before and after the chronic stress manipulation. The Open Field Test is a canonical behavioural test commonly used in rodent research as a measure of stress, based on the natural tendency of mice to exhibit thigmotaxis (remaining close to the periphery or walls of a novel surrounding) when stressed [23]. A testing arena (Plexiglass, 28 cm x 17 cm x 12 cm) was divided into a circular “Central Zone” (radius 6.35 cm, area = 127 cm^2^) and “Exterior Zones” (remaining area = 349 cm^2^). The Exterior Zone comprises a larger area than the Central Zone, but this is standard in protocols for the Open Field Test [48], and still allows for a reliable behavioural assessment of stress. Mice were placed in the centre of the arena and the total time spent with all four feet within the Central Zone over a 5-minute session was recorded.

#### Urine Corticosterone ELISA

An elevation in corticosterone levels is a natural and reliable response to the experience of stress in mice [1]. Previous studies have used urine corticosterone ELISAs to measure stress in mice [53, 34]. Urine samples were collected from mice using Fitchett’s method [19] immediately following the Open Field Test during both the pre-stress and post-stress sessions. Mice were transferred to an empty Plexiglass cage similar to their regular housing cages and allowed to roam freely. Samples were collected using a micropipette after the mice urinated and pooled for each mouse to collect approximately 100 *µ*L of urine for each mouse. Samples were stored at -80°C immediately after collection.

To measure corticosterone levels, the samples were thawed to room temperature and diluted 20-fold with SBS diluent as per the assay manufacturer’s recommendation. ELISA assays were only performed for a subset of mice (*n* = 20) in the study - 10 socially-housed mice and 10 singly-housed mice. Corticosterone concentrations in the urine samples were determined using corticosterone ELISA kits according to manufacturer’s instructions (Corticosterone ELISA Kit, AssayPro LLC, St. Charles, MO), with triplicates for each urine sample. Absorbances for samples were measured at 450 nm using the SpectraMax 190 Microplate Reader (Molecular Devices, San Jose, CA). Since the urine corticosterone ELISA is a competitive ELISA, absorbances were transformed into % Bound (*B/B*_0_) for all samples using Maximum Binding (*B*_0_) controls. Urine corticosterone concentrations (ng/mL) were determined from these transformed absorbances using 4-Parameter Logistic Regression for standard curve-fitting. Transformations and curve fitting were conducted using a web-based ELISA analysis tool at www.elisaanalysis.com.

#### Cost-Benefit Conflict (CBC) Task

All mice were tested on the CBC task twice, one day before and one day after the chronic stress manipulation. As during Pretraining, a T-maze was fitted with food wells and clip-on lamps. At the beginning of each trial, the food well in one arm was filled with pure chocolate milk (a high reward) paired with a very bright (1600 lumens) light (a high risk). The other arm had dilute (80% v/v) chocolate milk (low reward) and a dim (100 lumens) light (low risk; Figure 1). For the CBC task, each mouse underwent ten forced-choice training trials followed by 20 free-choice experimental trials.

**Figure 1:**
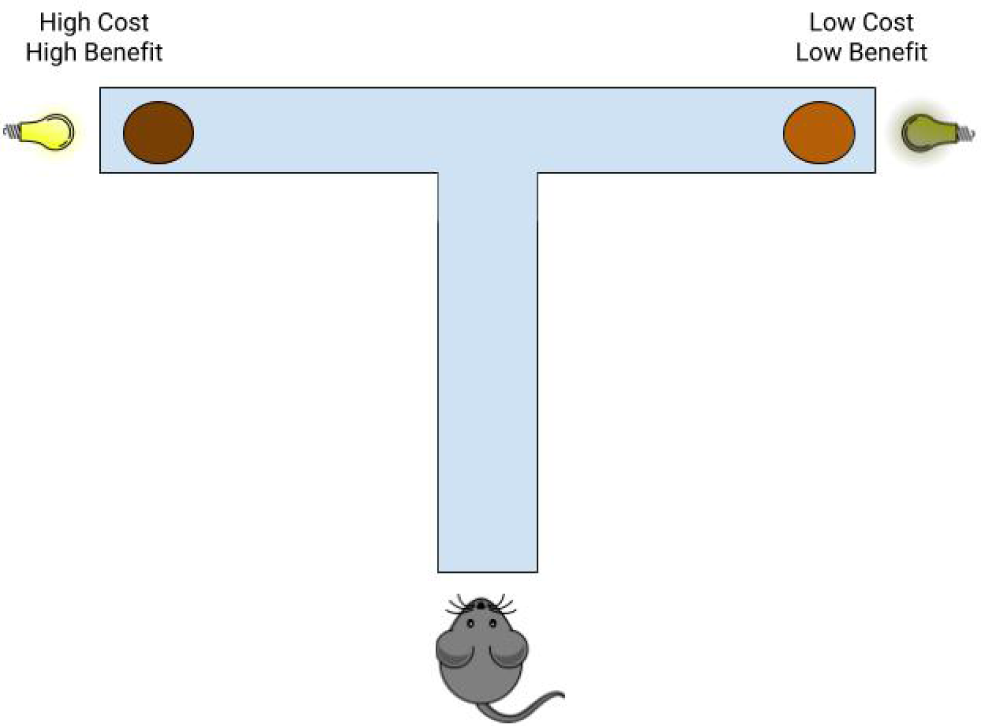
Schematic representation of the Cost-Benefit Conflict (CBC) task. A T-maze was used to assess decision-making in mice using the CBC task, in which mice chose between a high-risk/high-reward alternative (pure chocolate milk paired with bright light) and a low-risk/low-reward alternative (dilute chocolate milk paired with dim light).

##### Training Trials

In the ten training trials, the high-reward/high-risk and low-reward/low-risk alternatives were presented. However, one arm of the maze was blocked off using a guillotine door, so that the mice were forced to explore the only open arm. The arm that was open to explore was alternated between each training trial.

##### Experimental Trials

For the 20 free-choice experimental trials, mice were released onto the stem of the T-maze with both arms of the maze were open for exploration. A mouse was considered to have made a “decision” when it drank at least some chocolate milk from the food well of an arm. No decision was recorded if a mouse chose not to drink from either food well after 2 minutes on the maze. Data from the 20 experimental trials for each mouse were averaged to find % of decisions in which a mouse chose the high-risk/high-reward alternative.

#### Repeated Immobilisation Protocol

Mice were immobilised for one hour each day for seven consecutive days. They were immobilised using mouse restraint bags (Animal Identification and Marking Systems, Inc., Catalogue Number 89066-344). These restraint bags were small enough to significantly limit the animals’ mobility but large enough for the animals to breathe freely. Mice were placed on a flat surface and coaxed into entering the wide end of the bags, after which the tail end was tied with a zip-tie to prevent escape. Mice that were immobilised in this manner were placed on a flat surface and monitored regularly to ensure free breathing.

### Statistical Analysis

Our study was a 2 (Housing Condition: Single or Group) x 2 (Stress Exposure: Pre- and Post-) mixed design. We had three separate dependent measures: urine corticosterone concentration, time spent in the centre of the open field, and % of high risk/high reward decisions. We performed a linear mixed effects analysis for each of the 3 measures using R 3.6.1. The fixed effects were Housing Condition and Stress Exposure, and to account for intrinsic performance differences between mice, all analyses also included a random effect of mouse. We used estimated marginal means to perform post-hoc tests and corrected for multiple comparisons using Tukey’s method.

## Results

### Repeated immobilisation reliably induced stress, but social housing decreased stress-induced anxiety in the open field test

Data from the Open Field Tests and urine corticosterone ELISA assays were analysed to determine whether repeated immobilisation was successful in inducing chronic stress in the mice and whether housing condition affected stress measures. For both the Open Field and ELISA data, we ran a linear mixed effects model with two factors: Housing Condition (Singly or Socially) and Stress Exposure (Pre- and Post-Stress), with random effect of mouse.

#### Open field tests

The mean time spent in the centre of the open field (“open field activity”) was the dependent measure, where greater open field activity would indicate lower stress levels. The results (Figure 2) showed a significant main effect of Stress Exposure with a large effect size, *F* (1,30) = 197.50, *p* < .001, 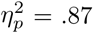, suggesting of stress protocol was effective. There was no significant main effect of Housing Condition, *F* (1,30) = 3.94, *p* = .06, 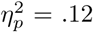. There was also a significant interaction of Stress Exposure and Housing Condition, *F* (1,30) = 13.44, *p* <.001, 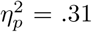, such that singly-housed mice showed a greater decrease in open field activity after repeated immobilisation than socially-housed mice (See Table 1 and 2 for group summary statistics and pairwise comparisons). The results show that our stress protocol was effective in inducing stress, and that it produced a greater change among singly-housed mice than socially-housed mice on a canonical behavioural test of stress.

**Table 1:** Group summary statistics and pairwise comparisons for our linear mixed effects models are presented here. Estimated Marginal Means and Standard Deviations for each group for all 3 dependent variables

| DV | Condition | Stress Exposure | Mean | Std Dev |
| --- | --- | --- | --- | --- |
| Open Field | Socially-housed | Pre-stress | 26.5 sec | 1.02 |
|  | Socially-housed | Post-stress | 20.3 sec | 1.02 |
|  | Singly-housed | Pre-stress | 26.1 sec | 1.02 |
|  | Singly-housed | Post-stress | 15.6 sec | 1.02 |
| ELISA | Socially-housed | Pre-stress | 3.19 (ng/mL) | 0.486 |
|  | Socially-housed | Post-stress | 8.44 (ng/mL) | 0.486 |
|  | Singly-housed | Pre-stress | 4.38 (ng/mL) | 0.486 |
|  | Singly-housed | Post-stress | 8.86 (ng/mL) | 0.486 |
| CBC | Socially-housed | Pre-stress | 36.5 (%) | 1.47 |
|  | Socially-housed | Post-stress | 52.9 (%) | 1.47 |
|  | Singly-housed | Pre-stress | 45.3 (%) | 1.47 |
|  | Singly-housed | Post-stress | 69.8 (%) | 1.47 |

**Table 2:** Group summary statistics and pairwise comparisons for our linear mixed effects models are presented here. Pairwise comparisons for each group for all 3 dependent variables.

| DV | Group | Comparison | $t(df)$ | p-value |
| --- | --- | --- | --- | --- |
| Open Field | Socially-housed | Pre-stress - Post-stress | $t(32.1) = 7.095$ | 0.000 |
| | Singly-housed | Pre-Stress - Post-stress | $t(32.1) = 12.11$ | 0.000 |
| | Pre-Stress | Socially - Singly | $t(45.8) = 0.223$ | 1.000 |
| | Post-Stress | Socially - Singly | $t(45.8) = 3.246$ | 0.013 |
| ELISA | Socially-housed | Pre-stress - Post-stress | $t(22.2) = -7.651$ | 0.000 |
| | Singly-housed | Pre-stress - Post-stress | $t(22.2) = -6.528$ | 0.000 |
| | Pre-stress | Socially - Singly | $t(44.4) = -1.735$ | 0.318 |
| | Post-stress | Socially - Singly | $t(44.4) = -0.611$ | 0.928 |
| CBC | Socially-housed | Pre-stress - Post-stress | $t(32.1) = -8.658$ | 0.000 |
| | Singly-housed | Pre-stress - Post-stress | $t(32.1) = -12.882$ | 0.000 |
| | Pre-stress | Socially - Singly | $t(62.5) = -4.243$ | 0.000 |
| | Post-stress | Socially - Singly | $t(62.5) = -8.096$ | 0.000 |

**Figure 2:**
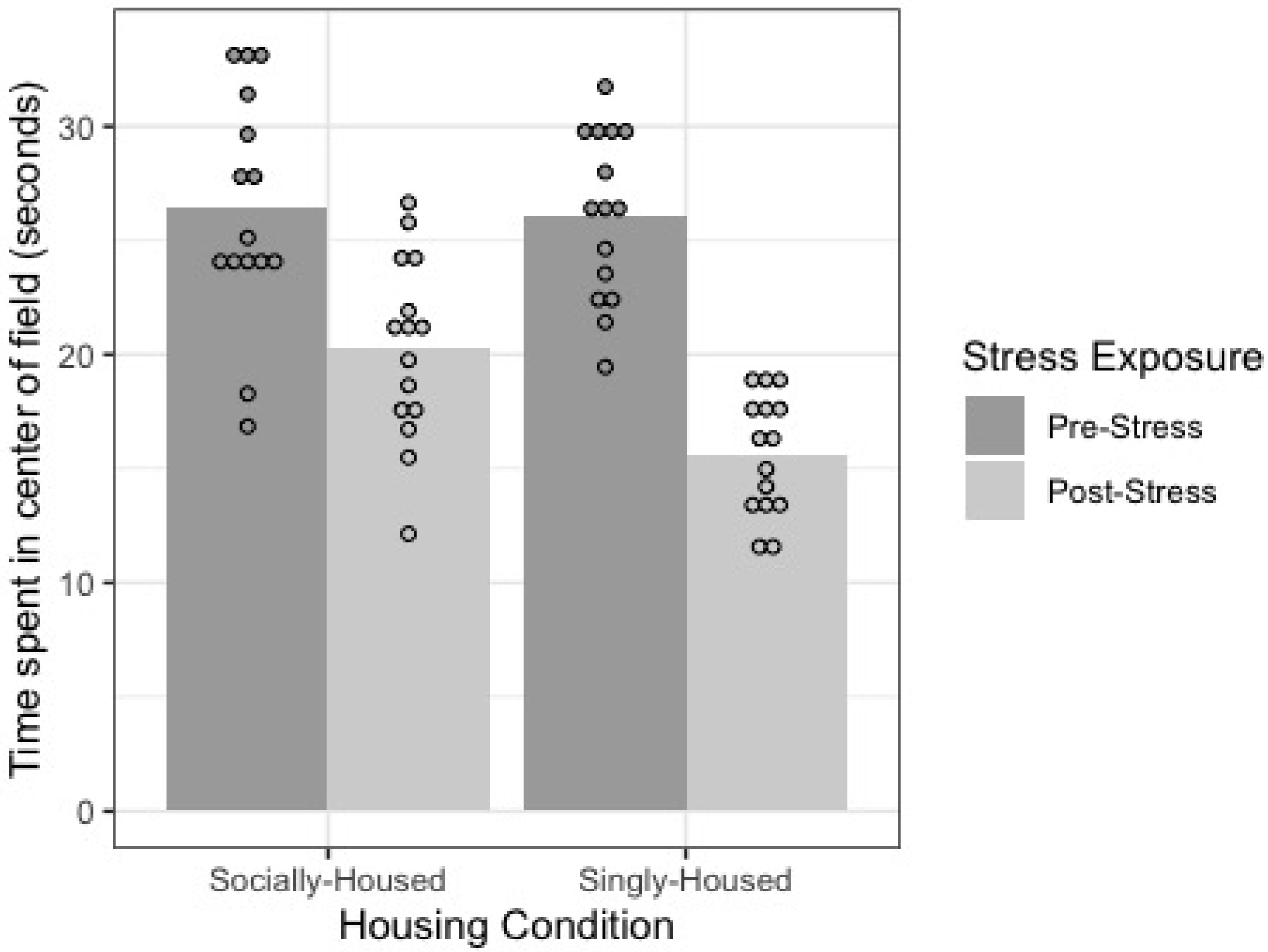
Performance of socially-housed (*n* = 15) and singly-housed mice (*n* = 15) on an Open Field Test before and after exposure to chronic stress. A decrease in mean time spent in the centre of the open field indicates an increase in behavioural stress levels. There was a significant interaction of Housing Condition and Stress Exposure, *F* (1,30) = 13.44, *p* <.001, 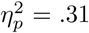, indicating that singly-housed mice showed a greater decrease in open field activity after exposure to chronic stress than socially-housed mice. Dots represent individual data points in each group, bars represent to mean for each group. See Table 2 for pairwise comparisons.

#### Urine corticosterone

Mean urine corticosterone concentration (ng/mL) was the dependent measure (Figure 3), where greater urine corticosterone levels would indicate higher stress levels. Results showed a significant main effect of Stress Exposure with a large effect size, *F* (1,40) = 111.69, *p* < .000, 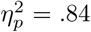. There was not a significant main effect of Housing Condition on urine corticosterone levels, *F* (1,40) = 3.06, *p* = 0.09, 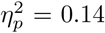, and no significant interaction *F* (1,40) = .70, *p* = .40, 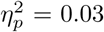. See Table 1 for group summary.

**Figure 3:**
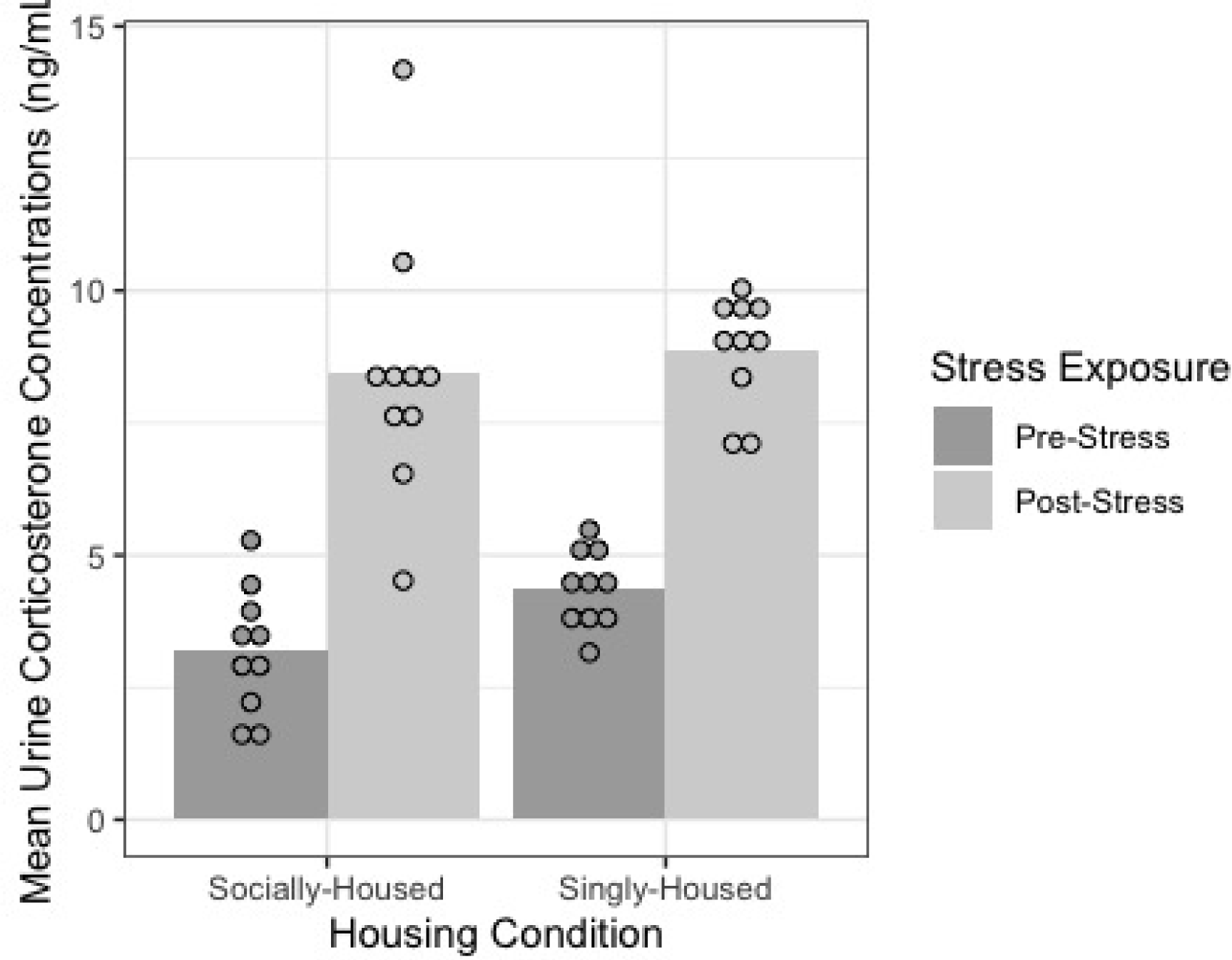
Mean urine corticosterone concentrations of socially-housed (*n* = 10) and singly-housed mice (*n* = 10) measured using urine corticosterone ELISA before and after exposure to chronic stress. An increase in mean urine corticosterone concentrations indicates an increase in physiological stress levels. There was no significant interaction of Housing Condition and Stress Exposure, *F* (1,40) = .70, *p* = .40, 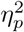, indicating that there was no significant difference in the increase in urine corticosterone levels between socially-housed and singly-housed mice. Dots represent individual data points in each group, bars represent to mean for each group. See Table 2 for pairwise comparisons.

Altogether, the significant main effects from the linear mixed effects models for the open field test and corticosterone ELISAs are sufficient to show that repeated immobilisation was successful in inducing stress as measured both physiologically and behaviourally. For the sake of thoroughness (and so that readers can see the complete data), we ran pairwise comparisons using the estimated marginal means (shown in Table 1) and using the Tukey’s method for multiple comparisons (Table 2).

### Socially-housed mice exhibited a smaller increase in high-risk decision-making after exposure to chronic stress than did singly-housed mice

We ran a linear mixed effects model, with two fixed factors Stress Exposure and Housing Condition. Mouse was a random factor. Mean % of decisions in which mice chose the high-risk/high-reward alternative was the dependent measure. There was a significant main effect of Stress Exposure on % of high-risk/high-reward decisions with a large effect size, *F* (1,28) = 248.53, *p* < .000, 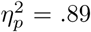, indicating that, in general, mice chose the high-risk/high-reward alternatives at a significantly higher rate after exposure to chronic stress than before. There was also a significant main effect of Housing Condition on % of high-risk/high-reward decisions, *F* (1,28) = 69.84, *p* < .000, 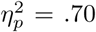, indicating that singly-housed mice chose the high-risk/high-reward alternatives at significantly higher rates than socially-housed mice. Finally, there was also a significant interaction between Stress Exposure and Housing Condition on the % of high-risk/high-reward decisions made by the mice, *F* (1,28) = 9.56, *p* = .004, 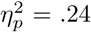. This significant interaction indicates that the increase in high-risk/high-reward choices after chronic stress differed between the socially- and singly-housed animals. Figure 4 and the means summarised in Table 1 and Table 2 show that the increase in % of high-risk choices by the socially-housed animals was smaller than the increase for singly-housed animals.

**Figure 4:**
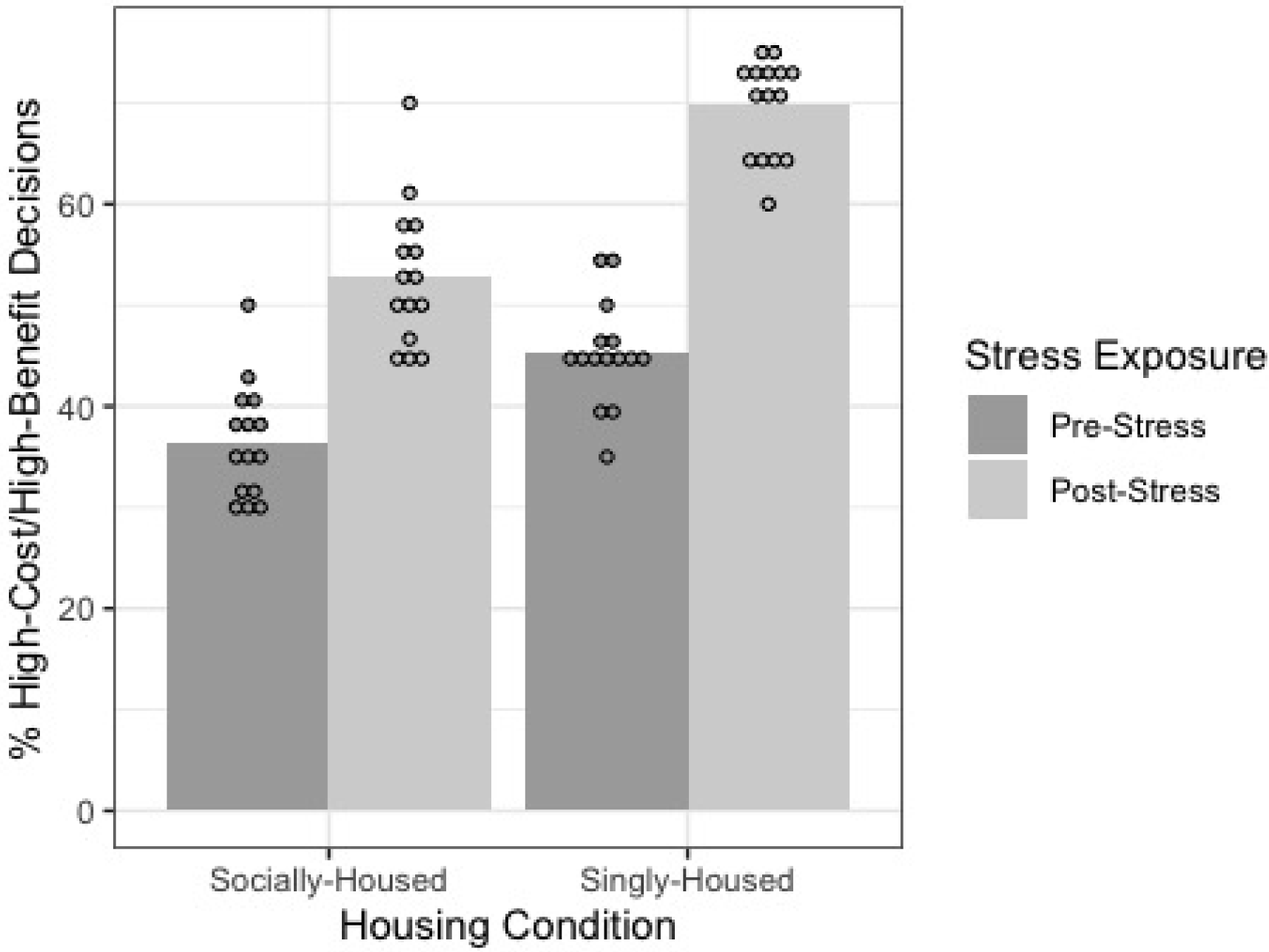
Performance of socially-housed (*n* = 15) and singly-housed mice (*n* = 15) on the Cost-Benefit Conflict task before and after exposure to chronic stress. An increase in % high-risk/high-reward decisions indicates an increase in preference for immediate payoffs (adaptive decision-making). There was a significant interaction of Housing Condition and Stress Exposure, *F* (1,28) = 9.56, *p* = .004, 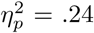, indicating that socially-housed mice showed a smaller increase in % high-risk/high-reward decisions than singly-housed mice after exposure to chronic stress. Dots represent individual data points in each group, bars represent to mean for each group. See Table 2 for pairwise comparisons.

## Discussion

This study investigated the role of social interaction in attenuating the effects of chronic stress on decision-making in mice. We hypothesised that mice would make more frequent high-risk/high-reward decisions after being chronically-stressed, and that this adaptive change in decision-making would be greater in mice that are housed singly than those housed in groups. Indeed, we found that, regardless of housing condition, mice showed an increased preference for the high-risk/high-reward alternative in the decision-making task after chronic stress exposure. This finding confirms the observations of [21]. Furthermore, the current study shows that socially-housed mice exhibited a smaller increase in high-risk/high-reward preferences compared to singly-housed mice. Critically, we have demonstrated that access to social interaction can have a mitigating influence on the effects of chronic stress.

In addition, the current study also confirms findings from previous research investigating chronic stress, social interaction, and cognition. In our study, mice were immediately assigned to singly-housed or socially-housed groups upon arrival of mice to our housing facilities. The mice were in these housing conditions for five days before pre-stress testing. Prior to the repeated immobilisation protocol, we observed differences between the housing groups on both the corticosterone measure and the CBC results (Table 2). This finding confirms previous literature which suggested that social isolation can have stress-like influences on the behaviour and physiology of rodents [37, 41]. Our study extends this knowledge by showing that social isolation can significantly alter decision-making.

One aspect of the study which limits generalisation of the results is that all the mice used in our study were adult male mice. We chose to use only male mice in our study to allow for comparison with previous studies that have investigated similar research questions such as [21], which also studied only male mice. However, studies do show that female mice exhibit different chronic stress responses than male mice [20], including differential behavioural and physiological reactivity [59, 46, 27]. Additionally, a recent study by [12] has suggested that female mice are more susceptible to nonsocial stressors compared to social stressors (such as isolation). In fact, social isolation may lead to opposite behavioural responses among male and female mice [40]. This further complicates our findings when attempting to generalise to female mice, since social interaction may influence stress responses differently in female mice than in male mice. We also used relatively young adult mice (about 8 weeks old), and certainly there are some age-related differences in behavioural and physiological responses to chronic stress among mice [52, 47]. Therefore, care must be taken when attempting to generalise the findings of the current study.

Despite this limitation, the current study makes strong contributions to our understanding of how chronic stress influences decision-making. Methodologically, the Open Field Test and urine corticosterone ELISA results confirmed that our short, 7-day repeated immobilisation protocol induced stress levels that led to measurable changes in both behaviour and physiology. These results thus affirm a much shorter stress-induction protocol than has been used in previous mouse studies of chronic stress effects [24, 21].

Additionally, our study demonstrates that chronic stress can cause changes in decision-making and lead to preferences for high-risk/high-reward decisions over lower risk alternatives that yield a lower reward. This could be due to a mechanism wherein chronically stressful experiences may create a mindset (or cognitive bias) that favours high rewards. Indeed, in conditions of high stress or low environmental predictability, a bias toward riskier immediate awards may in fact be adaptive for an organism. Further research needs to be conducted to examine the exact cognitive and physiological mechanisms for the effect we observed in this study. Moreover, studies can also directly investigate the adaptiveness of a high-risk/high-reward bias in stress environmental conditions using ecological methods.

Finally, we show that long-term social interactions decrease the effects of chronic stress, and thus, this research offers a model for mitigating the effects of chronic stress on cognition. Much more investigation is needed to understand the mechanisms and the limits of this effect. For instance, chronic stress due to repeated immobilisation may interact differently with social isolation than chronic stress due to social defeat [22], in which mice are repeatedly subjected to agonistic confrontations by a larger and more aggressive mouse leading to significant social stress. Since social defeat would arguably cause subsequent social interactions to be a negative affective experience for mice, social interaction may exacerbate, not mitigate, stress effects in mice that were chronically stressed through social defeat. Thus, while establishing a relationship between chronic stress and social isolation using one stress protocol, the current study also raises questions for future research about the influence of different types of stressors on this relationship.

Our findings have implications for human research into the effects of various stress-inducing experiences on cognition. In particular, results from mouse studies have been reliably extrapolated to studying conditions such as anxiety or depression in humans [11]. In fact, unpredictable chronic mild stress (UCMS), a well-established chronic stress protocol, has often been used in mouse studies to screen antidepressant drug candidates [38], underlining the value of mouse models in translational research into psychological disorders. The interrelationship between social interaction, chronic stress, and decision-making has significant implications for society, especially for professionals in high-stress jobs that often have to make decisions with wide-ranging consequences regularly, such as emergency medical professionals or finance consultants. Our findings suggest that healthy social interaction may play a crucial role in reducing the adaptive changes in decision-making that may be caused by chronic stress among these individuals.

One major insight from the current study is that chronic stress experience may cause changes in decision-making, which provides a new framework for understanding the factors that influence human cognition. Future research could explore what other stress-inducing experiences cause changes in decision-making. Are those changes domain-specific or domain-general? For example, does chronic social stress cause changes selectively to social cognition, or all cognitive processes generally? Another interesting avenue for research could be exploring systemic “minority stress” effects which have been observed among many marginalised population groups [35]. People living in poverty, LGBTQ+ individuals, and ethnic and racial minorities are some groups that experience both covert and overt discrimination in daily life, which can be thought of as chronic stress [16]. A growing body of health-related research suggests that this chronic marginalisation stress may negatively impact health and well-being in much the same way as more commonly studied forms of chronic stress [4, 6, 31, 57]. Findings from the current study support arguments that observed increases in preferences for immediate payoffs among these groups, often characterised as “irresponsible” behaviour inherent among marginalised groups, may in fact be an adaptive response to chronic stress. Furthermore, the current study offers evidence for the potential of social interaction to mitigate these changes in decision-making caused by chronic stress. With further research, findings from the current study may inform public health policies that combat chronic stress effects among marginalised populations.

The current study was the first to simultaneously examine the effects of chronic stress and social interaction on decision-making using a mouse model. We demonstrated that mice showed an increased preference for high-risk/high-reward alternatives after exposure to chronic stress regardless of housing condition. More importantly, socially-housed mice showed a smaller increase in proportion of high-risk/high-reward decisions after exposure to chronic stress when compared to singly-housed mice. Therefore, we found that chronic stress leads to an increase in high-risk decision-making in mice, an effect which is mitigated by social interaction. Through the current study, we offer a model for attenuating chronic stress effects on cognition among individuals. Our findings suggest that an increase in high-risk decision-making may be caused by chronic stress, and that social interaction may play a crucial role in combating these effects.

## Author Contributions

AMR conceived of the study, carried out the experiments, and wrote the paper. MTT aided in experimental design, data analysis, and manuscript preparation.

## Acknowledgments

The authors thank Daniel Thompson for support in purchasing animals and supplies, equipment setup, and lab resource management. AMR also thanks Dr. Peter Blair and Dr. Beth Mechlin for guidance on ELISA setups.

## Data Availability Statement

The datasets analysed for this study, as well as the code used for analysis, can be found in the repository named MudraRakshasa at https://github.com/michelletytong/MudraRakshasa.

